# The proteome of *Tetrasphaera elongata* is adapted to changing conditions in wastewater treatment plants

**DOI:** 10.1101/513143

**Authors:** Florian-Alexander Herbst, Morten S. Dueholm, Reinhard Wimmer, Per Halkjær Nielsen

## Abstract

The activated sludge in wastewater treatment plants (WWTP) designed for enhanced biological phosphorus removal (EBPR) experiences periodically changing nutrient and oxygen availability. *Tetrasphaera* is the most abundant genus in Danish WWTP and represents up to 20–30% of the activated sludge community based on 16S rRNA amplicon sequencing and quantitative fluorescence *in situ* hybridization analyses, although the genus is in low abundance in the influent wastewater. Here we investigated how *Tetrasphaera* can successfully out-compete most other microorganisms in such highly dynamic ecosystems. To achive this, we analyzed the physiological adaptations of the WWTP isolate *T. elongata* str. LP2 during an aerobic to anoxic shift by label-free quantitative proteomics and NMR-metabolomics. *Escherichia coli* was used as reference organism as it shares several metabolic capabilities and is regularly introduced to wastewater treatment plants, but without succeeding there. When compared to *E. coli*, only minor changes in the proteome of *T. elongata* were observed after the switch to anoxic conditions. This indicates that metabolic pathways for anaerobic energy harvest were already expressed during the aerobic growth. This allows continuous growth of *Tetrasphaera* immediately after the switch to anoxic conditions. Metabolomics furthermore revealed that the substrates provided were exploited far more efficiently by *Tetrasphaera* than by *E. coli*. These results suggest that *T. elongata* prospers in the dynamic WWTP environment due to adaptation to the changing environmental conditions.

**Significance of the study:** Members of the genus *Tetrasphaera* are widely distributed and highly abundant in most well-operating WWTPs with EBPR configuration. However, despite their high abundance *in situ*, little is known about their physiology and ecological role. Although the importance of *Tetrasphaera* in engineered wastewater treatment systems is slowly being recognized, additional information is needed to understand the full extent of functions the microorganisms have in many of the essential biological processes in the WWTP. Such information may improve available process models and ultimately lead to better wastewater treatment as well as resource recovery. This study supplies proteomic and metabolomic data on the aerobic/anoxic adaptation of *Tetrasphaera* and provides a hypothesis on how *Tetrasphaera* might compete in dynamic engineered systems.

## Introduction

Engineered environments allow for the enrichment of beneficial microorganisms by applying specific environmental conditions that benefit their function. In nature, the microorganisms may encounter changes in e.g. temperature or substrate as well as electron acceptor availability, and many of these changes follow irregular dynamic schedules. Microorganisms have developed different physiological adaptations to cope with such changes in carbon and electron donor/acceptor availability and they include use of storage compounds such as polyhydroxyalkanoate (PHA), lipids, glycogen, poly-phosphate (poly-P), elemental sulphur, and nitrate [1–3]. Very often adaptation also involves the expression of specific sets of genes [4], e.g., for acquisition and recycling pathways in case of substrate limitations [5], chaperones and proteases in case of heat [6], catalase or superoxide dismutase in case of oxidative stress [7], or expression of genes for alternative electron acceptors and fermentation in the absence of oxygen [8].

The adaptation of the microbial community to cope with dynamic conditions is exploited in modern wastewater treatment plants (WWTPs). By applying oxic/anoxic alternating phases and/or substrate rich and poor phases (feast-famine) as selective pressures, different microorganisms are enriched that efficiently remove carbon (C), nitrogen (N), and phosphorus (P) from domestic and industrial wastewater [2]. N is removed efficiently by nitrification/denitrification (oxic/anoxic) cycles which convert surplus N to inert N_2_ or biomass [9]. P is removed by enhanced biological phosphorus removal (EBPR) by including an anaerobic tank in addition to tanks for nitrification/denitrification and these plants can reach a high effluent water quality in respect to P with no or minimal chemical supplementation. In EBPR plants, poly-P accumulating organisms (PAOs) are enriched [10] and they store excess amounts of intracellular poly-P [2]. The general model assumes that poly-P is used for energy production during an anaerobic phase to take up carbon substrates. These are then stored as polyhydroxyalkanoates (PHAs) which are later oxidized aerobically in carbon-limited oxic phases for growth and poly-P regeneration. Over time P is depleted from the wastewater as the poly-P production is overall net-positive, and additional P is also incorporated into the biomass. Both can be removed as excess sludge and P can be recovered as raw material, e.g., before [11] or after anaerobic digestion [12].

The general EBPR model is primarily based on studies on the betaproteobacterial *Candidatus* Accumulibacter and the uptake of volatile fatty acids, mainly acetate, during an anaerobic phase, storage as PHA, and later oxidation in the aerobic phase [13,14]. Recently, another mechanism was revealed in PAOs belonging to the actinobacterial genus *Tetrasphaera*. Its members carry out aerobic heterotrophic growth and nitrate respiration, but they can also ferment. All these processes are important for a well-working EBPR process. Surprisingly, *Tetrasphaera* are not able to produce and store PHAs [15]. Instead, experiments using *T. elongata* str. Lp2 under dynamic anaerobic/aerobic conditions showed that they could accumulate free amino acids such as glycine for later oxidation in the aerobic phase [16]. The general model is undoubtedly appropriate for *Ca.* Accumulibacter and may explain many lab-scale and full-scale observations. However, microbial community analyses by fluorescence *in situ* hybridization (FISH) and 16S rRNA gene amplicon sequencing indicate that *Ca.* Accumulibacter accounts for only a small fraction of the PAO population in many EBPR plants and that *Tetrasphaera* is more abundant, occasionally reaching 20-30% of the biomass [17–19].

The aim of this study was to understand how *Tetrasphaera* can successfully compete in highly dynamic EBPR plants. Therefore, the proteome and extracellular metabolome (exometabolome) of aerobically grown *T. elongata* str. Lp2 cells before and after a 3 h anoxic phase were analyzed. *E. coli* str. K-12 was treated in the same way to serve as reference. *E. coli* was chosen as it can grow anaerobically by fermentation and nitrate reduction and it is being introduced in high amounts by the incoming wastewater without constituting any major fraction in WWTPs [18]. Also, *E. coli* is generally very versatile and can survive under many different conditions. This specific strain is further one of the best described microorganisms and good annotations as well as literature are available. As for most bacteria, the expression of the necessary genes for anaerobic energy harvest is strictly hierarchically controlled [8,20] and adaptation needs some time. The acquired data indicated that this is not the case for *T. elongata* and that it does indeed show a high level of metabolic robustness and readiness. This, together with poly-P as energy storage compound and a metabolic diversity, could partly explain continuous growth and successful competition in the dynamic WWTP environment.

## Materials and Methods

### Cultivation and sampling

*T. elongata* str. Lp2 and *E. coli* str. K-12 were cultivated in modified R2A (minimal) medium [16] to be comparable with previous studies. Inoculation was performed from liquid overnight cultures to an optical density (OD) at 600 nm of 0.01. Initial oxic cultivation was done in 50 ml medium within 250 ml conical flasks (25°C, 150rpm). For the 3 h of anoxic incubation, cultures were transferred to serum flasks, and oxygen was removed by repeatedly replacing the headspace with >99.9% pure N_2_. Cultivations were performed in quadruplicates, and whole cultures were sacrificed at the end of the aerobic or anaerobic phase. Growth was determined by measuring the change in OD at 600 nm of 1 ml culture broth in a cuvette and with total protein concentration (see below).

### Metabolomics & Proteomics

All metabolomic and proteomic samples were obtained and measured as four biological replicates. Extracellular metabolites were extracted and analyzed by 600 MHZ NMR as previously described [21]. In short, 15 ml of culture supernatant were lyophilized, rehydrated in 600 μl D2O with TSP as standard, adjusted to a pH of 7, and recorded as 1D-NOESY at 298.1 K on a BRUKER AVIII-600 MHz NMR spectrometer equipped with a 5 mm cryogenic inverse tripleresonance probe. NMR signals were identified and quantified using ChenomX and the known TSP concentration as reference.

For protein extraction, cells were lysed (in 1 % sodium deoxycholate, 50 mM triethylammonium bicarbonate) using the FastPrep-96 Instrument (MP Biomedicals) for 1 min at 1600 rpm and the All FastDNA-96 (MP Biomedicals) lysis matrix. Protein concentrations were assessed by BCA method in triplicates, and approximately 20 μg of protein was subjected to a polyvinylidene fluoride membrane-based proteomic sample preparation [22]. The protocol was adjusted to 20 μg of protein, and the membranes were washed twice with 66% acetonitrile before equilibration with an 8 M urea solution. Tryptic peptides were measured by nLC-MS/MS (Ultimate 3000 coupled to a Q Exactive, Thermo Fisher Scientific, Waltham, USA) applying a 3 h method (∼140 min elution window). Details can be found elsewhere [21]. Mass spectra were analyzed by MaxQuant (v. 1.5.3.30) [23] as previously described [21], but with up to 4 allowed missed cleavages. Organism-specific protein databases in FASTA file format were obtained from UniProt [24]. Proteins were kept if they could be quantified (at least 2 peptides) in at least three replicates in one condition (aerobic or anaerobic). Label-free quantification (LFQ) values were used to compare the relative changes. The abundance data was log2 transformed and missing values were replaced by imputation from the normal distribution before statistical analysis. Significant changes in abundance were identified by t-test (two-tailed, permutation-based correction, 250 randomizations, FDR < 5%) in Perseus [25]. Plots were created using R [26] and the ggplot2 package [27]. The mass spectrometry proteomics data have been deposited to the ProteomeXchange Consortium (http://www.proteomexchange.org/) via the PRIDE partner repository [28] with the dataset identifier PXD005211.

## Results and Discussion

The genus *Tetrasphaera* is repeatedly observed as the most abundant genus in many EBPR plants [17–19]. Since dynamic conditions with oxic/anoxic changes of roughly 3-4 h duration are fundamental to these plants, the reaction to oxygen deprivation under controlled conditions was investigated using *T. elongata* str. Lp2. This species has been used as the model organism for this clade of PAOs as it has a relatively high similarity on 16S rRNA gene level to *in situ* abundant phylotypes [17].

### Efficient growth under anoxic conditation

Pure cultures of *E. coli* and *T. elongata* were grown in modified R2A medium (without starch) and they showed, as expected, different growth patterns. *T. elongata* grew considerably slower than *E. coli* and required approx. three times as long time to reach a similar amount of biomass before the start of the anaerobic phase (24 vs. 7 hours). During the anoxic growth period, relative and absolute growth were at least as high for *T. elongata* as for *E. coli* as determined by biomass differences as assessed by the change in OD and total protein concentration (Tab. 1).

**Table 1.** Growth of *E. coli* and *T. elongata* after shift to anoxic conditions. Growth was determined by optical density (OD) at 600 nm and protein concentration after cell lysis. The mean and standard deviation of four measurement is presented.

|  | OD<sub>600nm</sub> |  | Protein concentration |  |
| --- | --- | --- | --- | --- |
|  | *E. coli* | *T. elongata* | *E. coli* | *T. elongata* |
| Aerobic phase (t = 0 hr) | 0.183 ± 0.003 | 0.174 ± 0.026 | 107.8 ± 0.8 µg/mL | 78.4 ± 3.1 µg/mL |
| Anoxic phase (t = 3 hr) | 0.214 ± 0.009 | 0.209 ± 0.032 | 114.0 ± 5.1 µg/mL | 88.7 ± 1.8 µg/mL |
| Absolute increase | 0.031 ± 0.009 | 0.036 ± 0.015 | 6.2 ± 5.3 µg/µL | 10.3 ± 3.8 µg/µL |
| Relative increase | 16.8 ± 4.8 % | 20.7 ± 9.2 % | 5.8 ± 5.0 % | 13.2 ± 5.3 % |

### Cost-effective use of substrates

To obtain detailed information of the substrate usage and the production of fermentation products, the medium was analyzed by NMR at different growth stages (Fig. 1). *T. elongata* was found to use considerably less sugars and amino acids for growth compared to *E. coli*. Surprisingly, *T. elongata* first depleted trehalose in the medium and subsequently exploited glucose and aspartate. Other amino acids were not visibly used as substrates, although *T. elongata* can metabolize all the measured amino acids in glucose-free medium. Previous genomic analyses suggested that *T. elongata* might produce lactate as fermentation product [15]. This could not be observed in this study and emphasizes the need for methods that provide direct information about phenotypic behaviour, such as metabolomics. Under the conditions applied, the only fermentation product observed in significant amounts was succinate. Both *E. coli* and *T. elongata* showed increased succinate production during anaerobic incubation.

**Figure 1.**
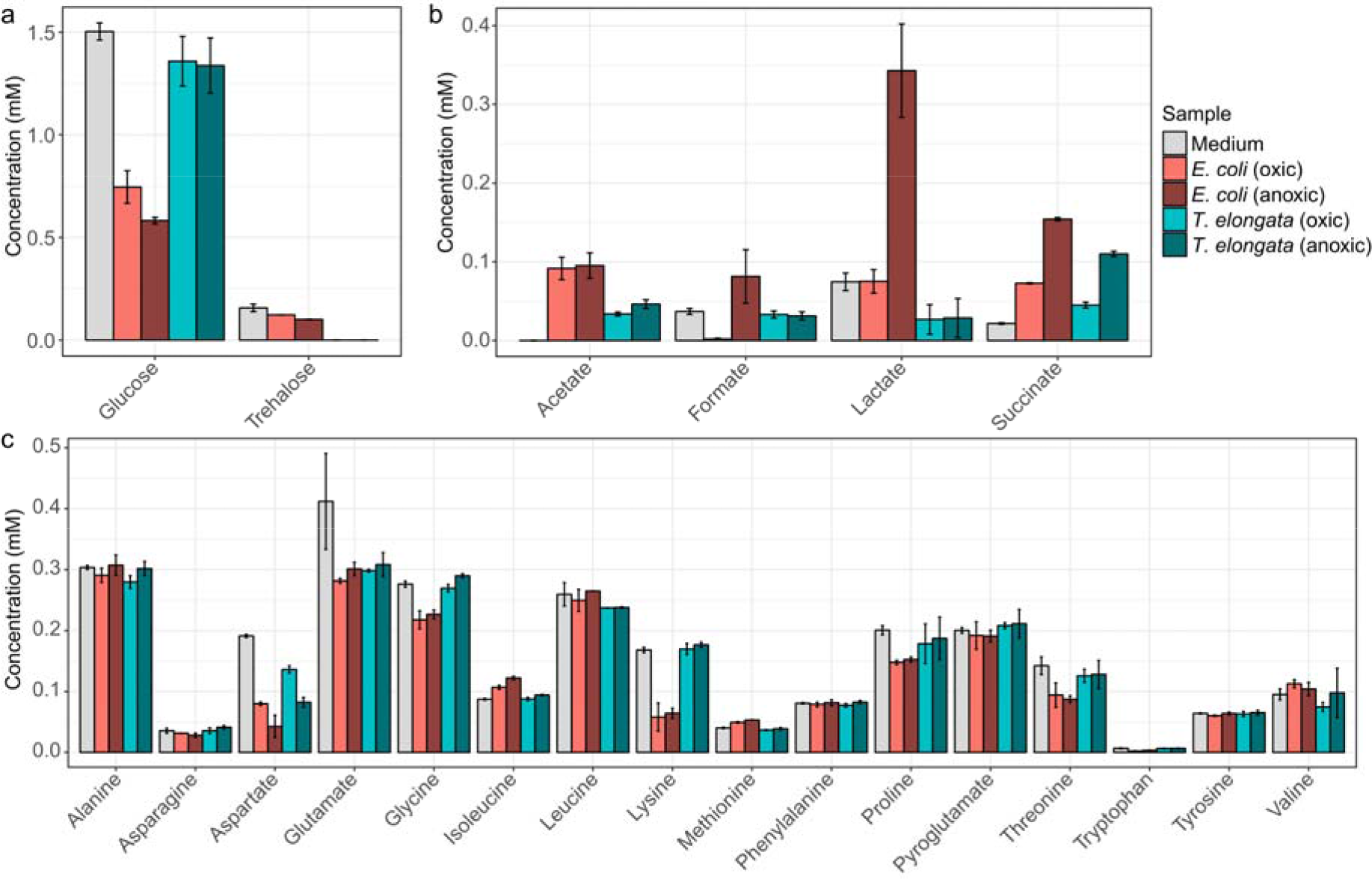
NMR-quantified exometabolites. (a) carbohydrates, (b) fermentative products, and (c) amino acids in the initial medium as well as for the end of the aerobic and anoxic cultivations of *E. coli* and *T. elongata*. Error bars represent standard deviation of quantified metabolite levels (n = 4).

Substrates were not a limiting factor in the experiments (Fig. 1), although provided in low concentrations, as they are in WWTPs and thus the oxic/anoxic switch can be seen as the primary environmental variable.

### Minor changes in the proteomes upon shift to anoxic conditions

It was possible to quantify 1318 and 1224 proteins or 42.7% and 28.5% of the theoretical proteome for *T. elongata* and *E. coli*, respectively (up to 1703 identified proteins, see Dataset1.xlsx). Out of these, 228 and 171 were significantly altered in abundance (FDR 5%), but with higher maximum fold changes in *E. coli* (Fig. 2a). Classification of the differentially expressed proteins based on COG class IDs [29] revealed considerable larger changes in the abundance of proteins related with energy production and conversion, amino acid metabolism and transport, carbohydrate metabolism and transport, and inorganic ion transport and metabolism for *E. coli* compared to for *T. elongata* (Fig. 2b).

**Figure 2.**
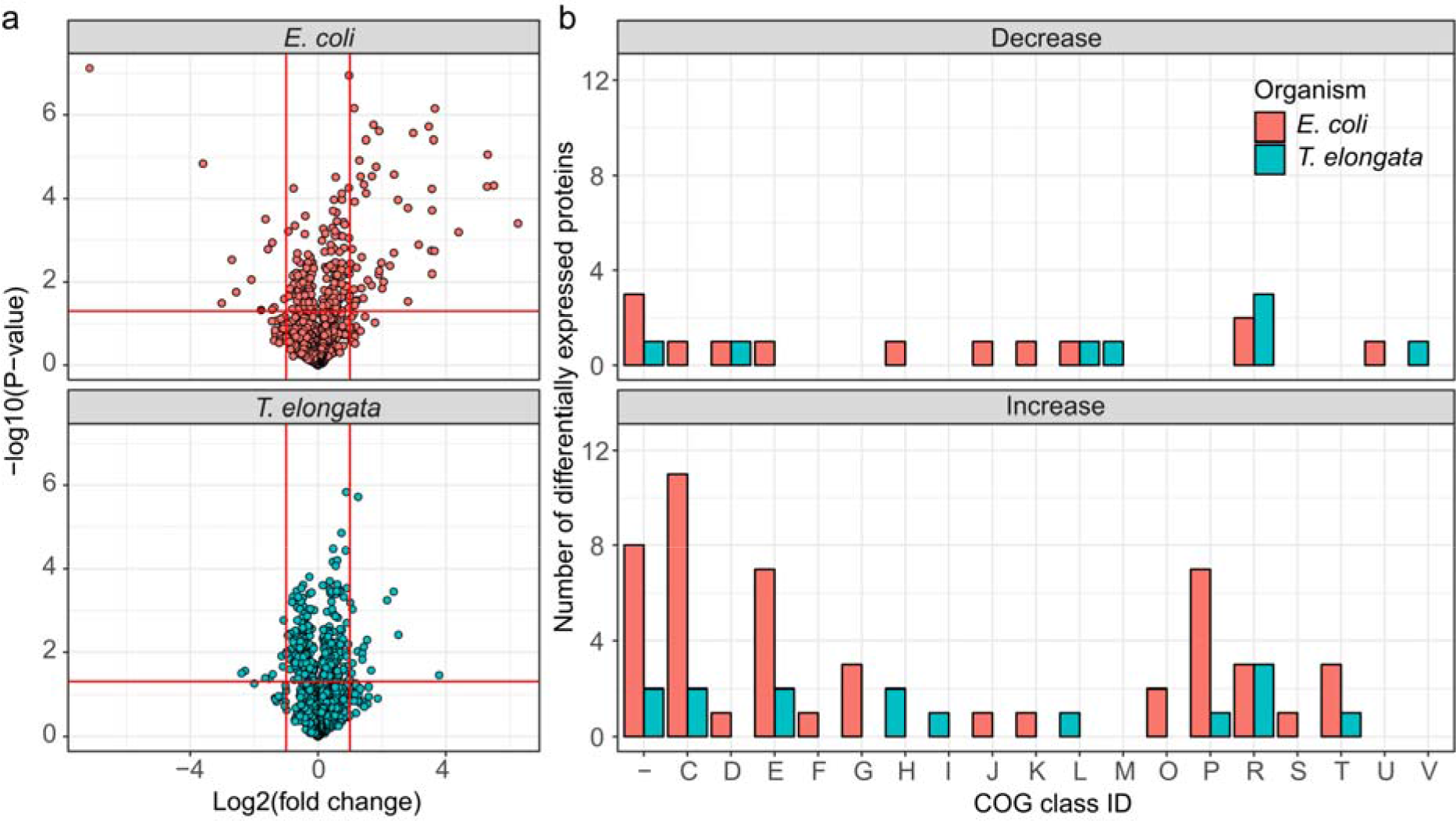
Overview of differentially expressed proteins for the end of the aerobic and anoxic cultivations of *E. coli* and *T. elongata*. (a) Volcano plot for quantified (after imputation) proteins from *T. elongata* and *E. coli*. Significance testing was corrected for multiple hypotheses testing at an FDR of 5%. Vertical red lines indicate a twofold change in abundance. The horizontal line reflects a p-value threshold of 0.05. (b) Classification of differentially expressed proteins based on COG functional classes. One-letter abbreviations for the functional categories: C, energy production and conversion; D, cell division and chromosome partitioning; E, amino acid metabolism and transport; F, nucleotide metabolism and transport; G, carbohydrate metabolism and transport; H, coenzyme metabolism; I, lipid metabolism; J, translation, including ribosome structure and biogenesis; K, transcription; L, replication, recombination and repair; M, cell wall structure and biogenesis and outer membrane; N, secretion, motility and chemotaxis; O, molecular chaperones and related functions; P, inorganic ion transport and metabolism; R, general functional prediction only; T, signal transduction; S and “-“, no functional prediction.

Looking at specific changes for proteins involved in energy production and stress response, the proteomic similarities between the two organisms were primarily restricted to an aspartate ammonia-lyase (AspA) and, interestingly, to an ATP-dependent RNA helicase, which was the most significantly down-regulated protein in both proteomes: DeaD and HelY in *E. coli* and *T. elongata*, respectively. These RNA helicases are required to adapt to different environmental situations [30]. AspA provides fumarate, presumably for reduction to succinate, from aspartate. For *E. coli*, it has been shown that AspA is part of the NarL regulon [20]. These proteomic adaptations fit the observed depletion of aspartate in the medium. The aspartate-succinate conversion has been reported for several bacteria [31–33] and yeast [34] and might be a general anaerobic adaptation and be preferred to the phosphoenolpyruvate to oxaloacetate conversion (slight, but significant up-regulation in *T. elongata* of PckG). Although succinate was undoubtedly the dominant fermentative product for *T. elongata*, no fumarate reductase [35] for fermentation is annotated in its genome. For *B. subtilis* it was shown that the ‘aerobic’ succinate dehydrogenase could fulfil this role [36] and *T. elongata* might do likewise. No other fermentation products were of importance to *T. elongata*. Although it depleted lactate under aerobic conditions, it did not produce any during anaerobic incubation. However, the lactate dehydrogenase (Ldh) could be identified in both conditions. *E. coli* did produce high amounts of lactate in the absence of oxygen (and other terminal electron acceptors), and the necessary lactate dehydrogenase (LdhA) was significantly more abundant. This assumingly led to additional acidic stress which was compensated by the expression of a lysine decarboxylase (CadA) which is regarded as a major acid stress enzyme [37]. Apart from the increased abundance of AspA, the hierarchical control of anaerobic gene expression [20,38] was evident only for *E. coli* (Fig. 3). Enzymes for fermentation or nitrate reduction were identified already under aerobic conditions for *T. elongata* and did not change significantly in their relative abundance during anaerobic incubation. In contrast to this, enzymes for nitrate reduction or fermentation were not identified before or showed clear up-regulation during anaerobic incubation for *E. coli* (Fig. 3). The presence of the anaerobic metabolic machinery for energy production in *T. elongata* explains their ability to ignore the anaerobic shock and keep a steady growth. Under relatively stable conditions like in the human gut this will probably be a costly disadvantage, but in dynamic systems like wastewater treatment plants with regular environmental changes, this might be a major cost-saving advantage.

**Figure 3.**
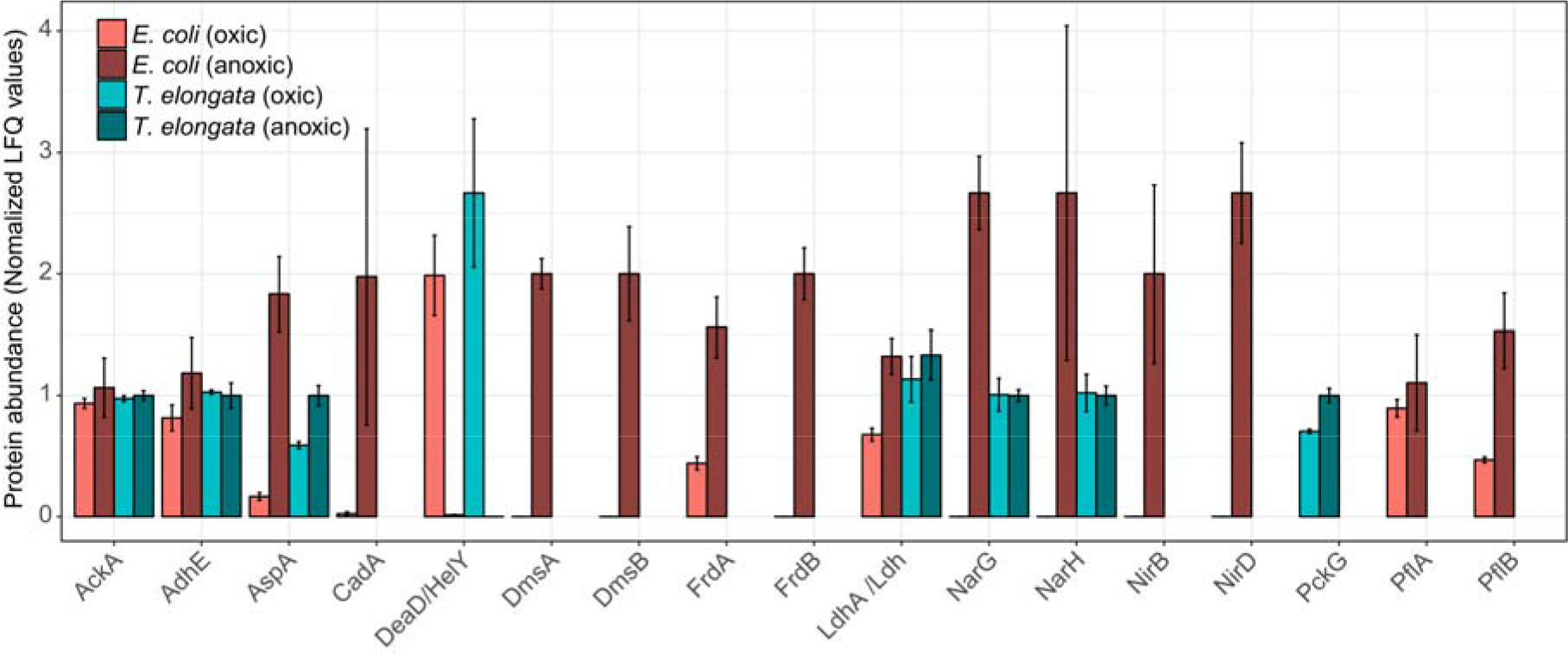
Normalized abundance of selected proteins for the end of the aerobic and anoxic cultivations of *E. coli* and *T. elongata*. The estimated abundance of each protein in each organims based on LFQ values was nomalized based on the average abundance of that protein across samples. Error bars represent standard deviation of protein abundance levels (n = 4).

### PAO-metabolism in T. elongata

As a PAO, poly-P accumulation and degradation in *T. elongata* are of special interest. One central protein, which might be indicative of a PAO physiology, is the low-affinity Pit phosphate transporter [39]. In the well-described PAO *Ca.* Accumulibacter, Pit seems to drive VFA uptake in the anaerobic phase [40]. In accordance with previously described observations, the abundance of Pit did not show any significant change between oxic and anaerobic conditions for *T. elongata*. Neither did the high-affinity Pst system. A small (∼1.3 fold), but significant, upregulation in the anaerobic phase could be observed for the polyphosphate kinase Ppk2. In contrast to Ppk1, which is widely conserved in bacteria [41] and did not show any significant change, Ppk2 regenerates GTP [42] as well as ATP [43] by utilizing poly-P, potentially providing additional energy during anaerobic conditions. Another poly-P utilizing enzyme which was slightly, but significantly, more abundant was the polyphosphate glucokinase PpgK. PpgK serves as an alternative hexokinase, which can phosphorylate glucose to glucose-6-phosphate by exploiting poly-P instead of ATP [44], thus, providing glucose-6-phosphate for glycolysis while preserving ATP. Apart from poly-P, glycogen is supposed to be the major element in *Tetraphaeras*’s PAO metabolism. Based on genomic and biochemical analyses, free amino acids and glycogen are the major C storage under anaerobic conditions [15,45]. A recent comprehensive *in situ* study using Raman spectroscopy could not verify the accumulation of glycogen in *Tetrasphaera* [46]. In this study, several putative glycogen related enzymes for synthesis (GlgB, GlgC, N0E1Q7, N0E176) and degradation (GlgX, GlgP) could be identified. Only N0E1Q7, a 1,4-alpha-glucan branching enzyme, showed a statistically significant, but very small (∼ 1.2 fold) change in abundance under aerobic conditions. This change was contrary to the assumption that glycogen synthesis is required during anaerobic conditions and carbon storage in *Tetrasphaera* remains a question of interest.

Overall, the data suggest that *T. elongata*, in contrast to *E. coli*, is pre-adapted to anaerobic conditions with the pathways necessary for energy production at the ready. If oxygen as a terminal electron acceptor is unavailable, *E. coli* reacts by first using alternative electron acceptors like nitrate or fumarate and ultimately relies on fermentation. As for most bacteria, the expression of the necessary genes is strictly hierarchically controlled [8,20] and adaptation needs time. The acquired data indicated that this is not the case for *T. elongata* and that it does indeed show a high level of metabolic robustness and readiness [47]. The theoretical advantage of being able to keep growing without major metabolic adaptations will be enhanced under *in situ* conditions as *T. elongata* can store excessive amounts of poly-P for rapid supplementation of energy or accumulation of carbon substrates under anaerobic conditions [16]. The proteomic and metabolomic data helps to complement the existing metabolic model [15] (Fig. 4) and gives an explanation for the abundance of *Tetrasphaera*. Although *Ca.* Accumulibacter and *Tetrasphaera* differ in central aspects of their metabolism and substrate preferences, they might follow a similar strategy. In that sense, no marked effects on relative protein abundances were observed across an EBPR cycle for *Ca.* Accumulibacter [48,49]. On the other hand, clear indications of regulation could be found on transcriptome level for *Ca.* Accumulibacter [50] which might not have been recognizable due to sample complexity and technical challenges. Unfortunately, transcription and translation do not always correlate [51] or are at least time-delayed [52] and conclusions can be challenging. Here, it was possible to observe the small but significant effects on proteome level which might not be observable in complex *in situ* experiments. A question that arises is, if the observations are entirely correct on the single-cell level or if community level adaptations are responsible. *T. elongata* has a more pronounced tendency to aggregate compared to *E. coli*. An advantage in WWTPs, but a disadvantage for cultivation in the laboratory. This study was performed at low cell-densities, vigorous shaking in large flasks with little culture volumen to reduce aggregation and its impact on physiology. Nevertheless, aggregates cannot be excluded which would lead to pre-adaptation of some *T. elongata* cells which enable constant growth in regularly changing environments. In favor of the results reflecting single-cell adaptations is that, e.g., the nitrate reductase (alpha and beta subunits) did not change in abundance which might be expected if all cells are now challenged with oxygen limitation (instead of a minor fraction that is already expressing the enzyme) as was observed for *E. coli*. *T. elongata*’s behavior is somewhat paradox. *T. elongata* is preventively expressing unnecessary genes (e.g., nitrate reduction) while showing an economical metabolism with relatively low growth rates and without strong reactions to environmental stimuli (at least oxygen limitation). Of course, predicting *in situ* responses from *in vitro* data must be done carefully, but the observations and the concluded hypothesis fit *in situ* observations and will guide future enrichment reactor and *in situ* studies.

**Figure 4.**
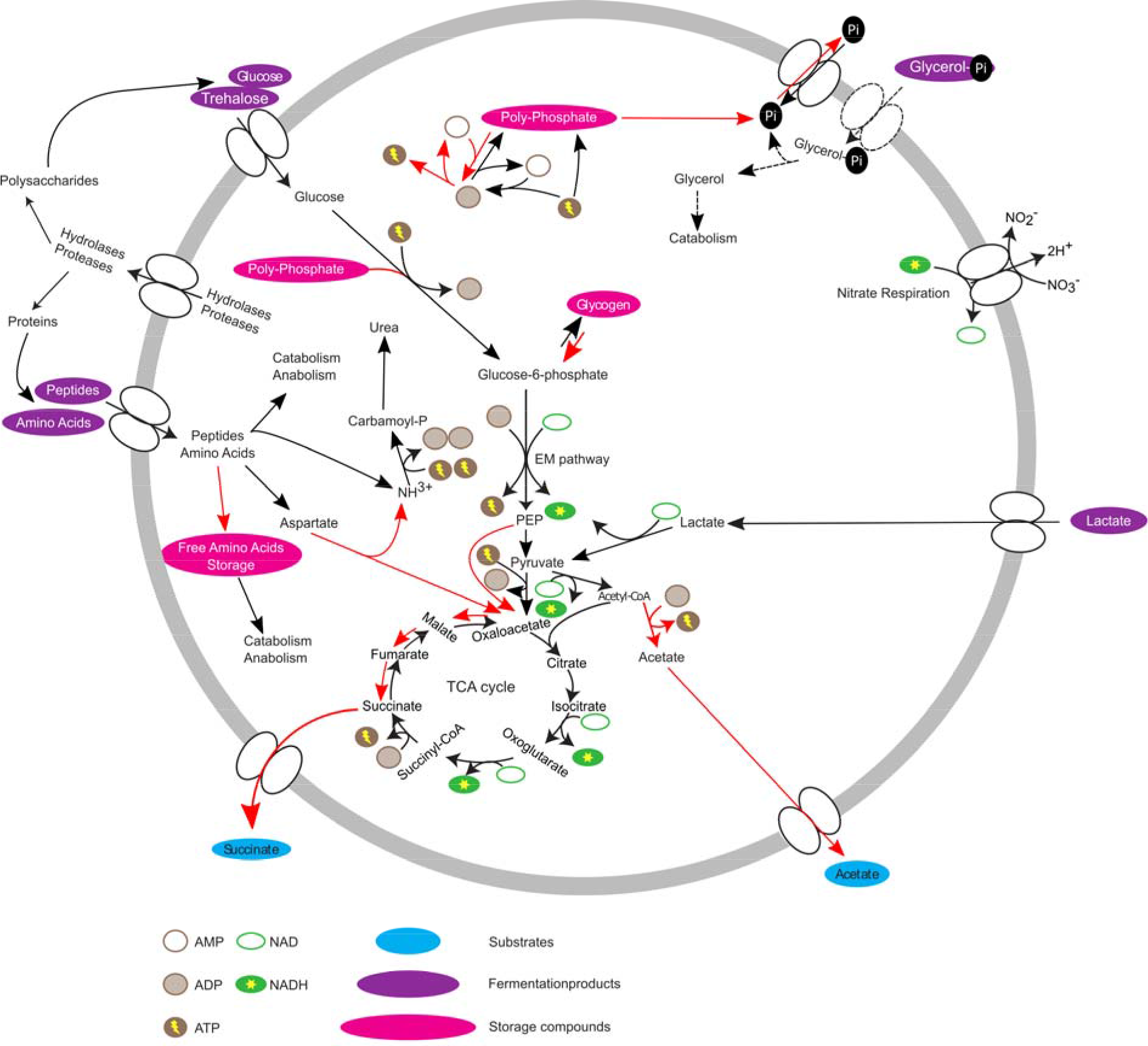
Updated metabolic model of *Tetrasphaera elongata* str. Lp2. Red arrows indicate increased activity under anaerobic conditions. Pink ovals are storage compounds, lilac ovals are substrates, and blue ovals are fermentation products.

## Concluding remarks

As expected, *E. coli* grew initially faster than *T. elongata*, but *T. elongata* grew consistently during the 3 h anaerobic switch. *T. elongata* behaved more economically to produce similar amounts of biomass. Furthermore, *T. elongata* mainly used succinate fermentation as an electron sink, whereas *E. coli* produced succinate, acetate, formate/CO_2_, and high amounts of lactate. The proteomic data lead to the identification of roughly 200 statistically significantly regulated proteins for both organisms, but the regulation in *T. elongata* resembled more a fine-tuning of already present pathways whereas *E. coli* underwent major rearrangements. Analyses of enriched pathways in *E. coli* showed a clear down-regulation of pathways necessary for translation and aerobic respiration as well as an up-regulation of anaerobic respiration, fermentation, and severe stress (probably due to lactic acid production) when challenged with oxygen limitation. At the same time, *T. elongata* already expressed necessary pathways under aerobic conditions, ignoring the classic hierarchical control of anaerobic gene expression and just fine-tuned its metabolism. While this strategy might not be suitable for many natural environments, it seems well suited for engineered habitats like WWTPs and their scheduled dynamics.

## Acknowledgments

This study was supported by the Danish Research Council for Independent Research (FNU), grant no. 4002-00455B. The NMR laboratory at Aalborg University is supported by the Obelske Family, SparNord, and Carlsberg Foundations. We thank the PRIDE team for providing data deposition support.

